# Apoptotic bodies from patients with Sjögren’s disease drive atypical memory B cell and macrophage activation and autoantibody production

**DOI:** 10.64898/2026.07.15.738605

**Authors:** Adrian Y.S. Lee, Bo Shi, Sriram Gummadi, Pamali Fonseka, Tuong-Khanh My Tu, Quan Thinh Le, Ching-Seng Ang, Tien K. Nguyen, Ming Wei Lin, Thanh Kha Phan, Joanne H. Reed

**Author notes:** Correspondence to: Dr Adrian Lee, Dr Kha Phan or A/Prof. Joanne Reed, Dr Kha Phan La Trobe Institute for Molecular Sciences Science Drive, Bundoora, Victoria, Australia 3083 E Dr Adrian Lee or A/Prof. Joanne Reed Westmead Institute for Medical Research 176 Hawkesbury Road, Westmead, New South Wales, Australia 2145 E. Equal senior authors (Thanh Kha Phan, Joanne H. Reed).

## Abstract

The impaired clearance of apoptotic cells has long been associated with systemic autoimmune diseases like Sjögren’s disease (SjD), which is characterised by autoantibodies targeting antigens on apoptotic cells. However, most studies analysing human samples have used patient-derived phagocytes and *in vitro* generated apoptotic cells, which do not reflect the heterogeneity of patient-derived apoptotic bodies (ApoBDs). Here, we molecularly and functionally compare ApoBDs derived from patients with SjD and healthy donors. ApoBDs were increased in the circulation in SjD and induced atypical memory B cell expansion, which was attenuated by blocking B cell receptor (BCR) and Toll-like receptors (TLR) 7-9 signalling. Co-cultures of ApoBDs and B cells drove autoantibody production. Macrophages engulfing SjD ApoBDs had a distinct inflammatory profile compared to macrophages engulfing healthy control ApoBDs. These findings define patient-derived ApoBDs as immunostimulatory substrates capable of driving innate and adaptive immune responses consistent with the pathogenic features observed in SjD, hence opening a new avenue of diagnostic and therapeutic development.

## Introduction

Sjögren’s disease (SjD) is a common systemic autoimmune disorder frequently manifesting as dryness (sicca) symptoms of the eyes and mouth, fatigue and arthralgias. The disease is underpinned by B cell hyperactivity, inflammation with organ lymphocytic infiltration, and autoantibodies targeting nuclear autoantigens (1, 2). The seminal discovery that the nuclear autoantigens most frequently targeted in SjD (Ro52, Ro60 and La) cluster at the surface of dying cells(3) led to the hypothesis that the induction and pathogenesis of SjD is linked to defective clearance of apoptotic cells.

Apoptosis, a major form of programmed cell death, is essential for physiological development and tissue homeostasis. Downstream of apoptosis induction, dying cells fragment into membrane-bound extracellular vesicles of 1-5 μm called apoptotic bodies (ApoBDs) to promote rapid clearance (4). ApoBD removal, also known as efferocytosis, is a critical housekeeping function of professional phagocytes such as macrophages (5) where immune quiescence is maintained through anti-inflammatory mediators such as interleukin (IL)-10 and transforming growth factor (TGF)-β (6, 7). Efferocytosis is generally so efficient that the presence of apoptotic cells in human tissue is rare, even though billions of cells are estimated to die daily (8). However, sites of autoimmune pathology such as salivary gland biopsies from patients with SjD reveal an increased frequency of apoptotic cells (9, 10). Moreover, patients with SjD, systemic lupus erythematosus (SLE) and rheumatoid arthritis showed an increased concentration of ApoBDs in plasma compared to healthy donors (11). However, to our knowledge, these patient-derived ApoBDs have not been molecularly or functionally characterised.

Defective efferocytosis in autoimmune disease was initially hypothesised to be caused by dysfunctional professional phagocytes. Monocyte-derived macrophages (MDM) from patients with SLE displayed impaired clearance of autologous dying peripheral blood mononuclear cells, compared to non-autoimmune donors (12) and SjD-derived monocytes and MDM showed impaired clearance of apoptotic cell lines (13, 14). However, patient autoantibodies were also shown to contribute to defective apoptotic cell clearance. The Ro60 and La autoantigens not only redistribute to the cell surface during apoptosis but become accessible for binding by extracellular IgG autoantibodies (15, 16), which directly inhibited efferocytosis by healthy donor phagocytes compared to control IgG (14, 17, 18).

Impaired efferocytosis is linked to inflammation as unremoved dead cells transition into a lytic state termed secondary necrosis leading to the release of danger-associated molecular patterns and additional autoantigens (19, 20). Nuclear autoantigens combine with anti-nuclear autoantibodies, form immune complexes and elicit inflammatory cytokine secretion, including TNF and IFN-α, from co-cultured phagocytes in a Toll-like receptor (TLR)-dependent manner (21–23). Furthermore, mouse models deficient in key phagocytic proteins, such as C1q (24) and MerTK (25), showed impaired efferocytosis, spontaneous autoantibody production and developed kidney or salivary gland inflammation analogous to SLE and SjD patients. These models implicate defective clearance of apoptotic cells as a causative factor of autoimmunity and not just a byproduct of it.

While defective efferocytosis in autoimmunity is established, clinical translation of these findings to disease monitoring or therapeutic intervention has been limited. This may be partially due to an over-reliance on *in vitro* generated apoptotic cells for human studies, which do not contain the asynchronous and heterogeneous composition of native patient-derived ApoBDs. Here, we molecularly and functionally characterise ApoBDs derived from patients with SjD. ApoBDs are increased in SjD plasma relative to healthy donors, likely caused by increased apoptosis and defective efferocytosis in SjD patients. ApoBDs induce atypical memory B cell expansion, *de novo* autoantibody production and macrophage activation. Nuclear contents and autoantibodies in the SjD ApoBDs work in tandem to mediate these effects via B cell receptor (BCR) and TLR-dependent signalling. Together, our study defines patient-derived ApoBDs as immunostimulatory substrates capable of driving some of the pathogenic features observed in SjD, highlighting new therapeutic targets.

## Results

### Circulating apoptotic bodies are increased in patients with Sjögren’s disease, attributable to defective efferocytosis

We first sought to quantify circulating ApoBDs in SjD patients’ plasma, a proxy indicator of impaired efferocytosis (26), using flow cytometry analysis of forward scatter, FLICA^+^ (indicative of cleaved caspase 3/7) and annexin V^+^ (indicative of phosphatidylserine exposure) (27, 28) (Figure 1a) and cleaved PANX1 (Supplementary Figure 1a). Compared to age-matched heathy controls, SjD patients showed ∼6-fold increase in ApoBD count (Figure 1a). The quantity of circulating ApoBDs, however, did not relate to the EULAR SjD disease activity index (ESSDAI) (Figure 1b). To define the cellular origins of ApoBDs, we characterised circulating ApoBDs using lineage–specific immune markers (Supplementary Figure 1b). Approximately one–third of ApoBDs were derived from leukocytes (Figure 1c), with T cells and natural killer (NK) cells the dominant source (Figure 1d), despite a consistently low level of *ex vivo* detected apoptosis in these two populations compared to other leukocyte populations (Figure 1e and Supplementary Figure 1c).

**Figure 1.**
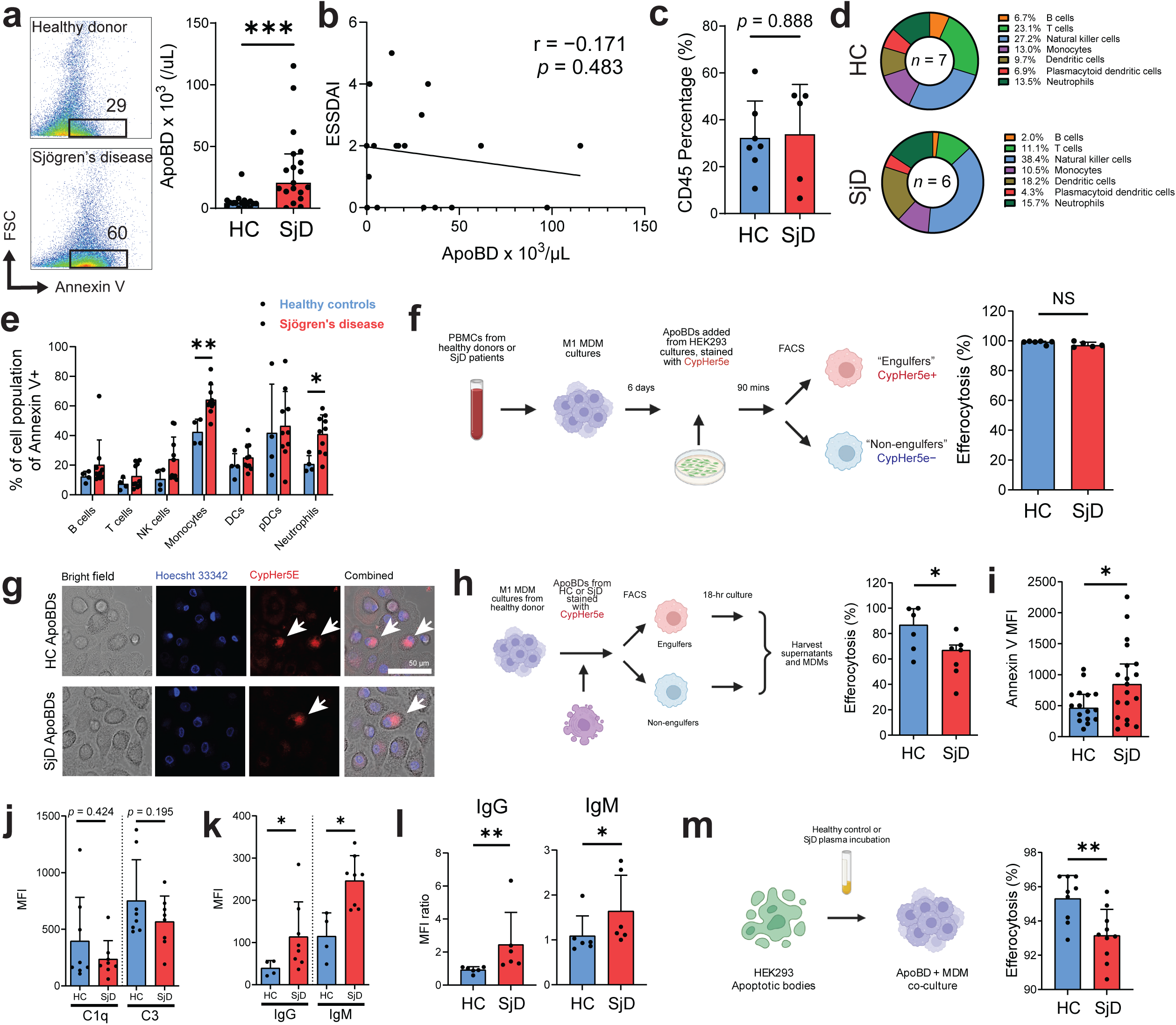
Circulating apoptotic bodies (ApoBDs) are increased in Sjögren’s disease (SjD) patients. **(a)** Flow cytometry quantification of circulating plasma ApoBDs defined as FLICA^+^, Annexin V^+^ and low forward scatter events. The plots (left panel) were pre-gated on FLICA^+^ events. ApoBDS were quantified from plasma obtained fresh from age- and sex-matched healthy donors (HCs) and SjD patients. The graph is pooled data from at least 10 independent experiments. **(b)** Scatterplot of ApoBD concentrations vs. EULAR SjD activity index (ESSDAI). **(c)** The percentage of ApoBDs positive for CD45 from HCs and SjD patients determined by flow cytometry. Data are pooled from 3 independent experiments. **(d)** Pie charts of the immune cell origins of ApoBDs in both HCs and SjD patients. Proportions are averaged across HCs and SjD groups and expressed as a percentage of total CD45^+^ cells. **(e)** Percent of annexin V^+^ cells for each peripheral blood immune cell population. Blood was processed within 1.5 hours of collection. Data are pooled from 6 independent experiments. *NK*, natural killer. *DCs*, dendritic cells. *pDCs*, plasmacytoid DCs. **(f)** Experimental design of monocyte-derived macrophage (MDM) cultures from HC and SjD peripheral blood co-cultured with ApoBDs derived from a HEK293 cell line at a ratio of 4:1 HEK293:MDM (left). Graph represents frequency of CypHer5E-positive MDMs per individual donor from 5 independent experiments. *NS*, not statistically significant by Mann-Whitney U test. **(g)** Micrograph images of MDM from HC peripheral blood mononuclear cells cultured with ApoBDs from HCs or SjD patients, labelled with CypHer5E dye. White arrows mark positive CypHer5E fluorescence in MDM cytoplasm confirming efferocytosis. Images are taken with 63× objective lens. Representative micrographs from two independent experiments of 2 HCs and 3 SjD patients. **(h)** ApoBDs derived from HCs or SjD patients were cultured with unrelated healthy donor MDMs and percent efferocytosis measured by flow cytometry. ApoBDs were cultured at a ratio of ApoBD:MDM 3:1. Graph represents pooled data from 6 independent experiments. **(i)** Phosphatidylserine on apoptotic bodies (ApoBDs) measured by annexin V mean fluorescence intensity (MFI) by flow cytometry. **(j)** Complement C1q and C3c deposition on ApoBDs. Complement MFIs were determined by flow cytometry. **(k)** Immunoglobulin deposition (IgG and IgM) on ApoBDs from HC and SjD by flow cytometry. **(l)** Binding of immunoglobulin to HEK293 ApoBDs measured by flow cytometry after incubating cells with diluted plasma 1:100 for 30 minutes at room temperature. MFIs ratios were calculated by dividing IgG or IgM MFIs for each donor plasma sample with a sample incubated with PBS only (background). Data are pooled from two independent experiments. **(m)** Efferocytosis of HEK293 ApoBDs pre-incubated with plasma from HCs or SjD. ApoBDs were cultured with healthy donor MDMs at a ratio of 3:1. Data is pooled from two independent experiments. For column graphs, each point represents an individual donor of ApoBDs or plasma. Columns represent mean with standard deviations. Schematic figures made with BioRender.

Consistent with previous observations (29), we saw elevated levels of apoptosis in SjD patients, particularly in monocytes and neutrophils (Figure 1d). The intrinsic capacity of healthy phagocytes is typically sufficient to accommodate increased apoptotic burden (30), making increased apoptosis alone unlikely to account for the increased circulating ApoBDs observed in SjD. Given that previous studies have shown reduced efferocytosis capacity of SjD-derived phagocytes (13, 14), we next compared uptake of apoptotic cell lines by MDMs from healthy donors and SjD. Surprisingly, no significant difference in the efferocytosis efficiency was found between SjD and healthy donor MDMs (Figure 1f). In contrast, when healthy MDMs were cultured with ApoBDs isolated from SjD or healthy donor plasma (Figure 1g), there was a significantly reduced uptake of SjD-derived ApoBDs (Figure 1h), suggesting that defective efferocytosis is attributable to the properties of apoptotic corpses rather than intrinsic phagocyte dysfunction.

To identify ApoBD-intrinsic features that may underlie the defective efferocytosis observed in SjD, we first assessed ApoBD surface phosphatidylserine exposure, a predominant “eat-me” signal for efferocytosis, using annexin V staining. SjD-derived ApoBDs exhibited increased surface phosphatidylserine compared with healthy control (Figure 1i), rendering deficient phosphatidylserine exposure unlikely to account for the observed impairment in efferocytosis. There were no significant differences in C1q or C3 complement deposition, which are also important “eat-me” signals and bridging molecules that aid efferocytosis (Figure 1j). Given that autoantibodies in SjD have been shown to impair clearance of apoptotic cells (14), we next examined surface-bound immunoglobulins (IgG, IgM) on circulating ApoBDs by flow cytometry. SjD-derived ApoBDs displayed significantly increased IgG and IgM relative to healthy controls (Figure 1k). To establish whether SjD serum-derived autoantibodies were contributing to reduced efferocytosis of SjD-derived ApoBD, we compared uptake of ApoBDs generated from HEK293 cells incubated with plasma from SjD patients or HCs. SjD patient plasma incubation caused greater IgG and IgM deposition on HEK293-derived ApoBDs (Figure 1l), which exhibited a mild reduction in efferocytic uptake compared to HEK293-ApoBDs incubated with healthy donor plasma (Figure 1m). Together, these findings indicate that SjD plasma contains autoantibodies that decorate ApoBDs and likely reduce efferocytosis relative to healthy donor plasma, thereby promoting an increase of apoptotic material in circulation.

### Sjögren’s disease-derived apoptotic bodies cause the expansion of atypical memory B cells and autoantibody secretion

Given that B cells in SjD patients are enriched for autoreactivity(31, 32) and atypical B cell subsets (32, 33), we hypothesised that ApoBDs may engage B cell receptors and directly influence B cell phenotype, subset composition and activation. To test this idea, primary human B cells from healthy donors were cultured with isolated ApoBDs from SjD or healthy donors *in vitro* for 48 hours and unbiased bulk proteomic profiling performed on B cells. Principal component analysis revealed distinct clustering of B cells that were treated with SjD ApoBDs from those treated with healthy control ApoBDs (Supplementary Figure 2a), consistent with the induction of a distinct SjD ApoBD-associated proteomic profile. Compared to healthy controls, SjD ApoBD co-cultures resulted in down-regulation of 43 and up-regulation of 1,116 proteins (Supplementary Figure 2b). Up-regulated proteins were enriched for metabolic and cellular processes consistent with heightened B cell activation (Supplementary Figure 2c). Gene ontology analysis of immune-related biological processes demonstrated significant enrichment of pathways involved in macroautophagy, antigen processing, B cell receptor signalling, and leukocyte migration in SjD ApoBD-treated B cells (Figure 2a).

**Figure 2.**
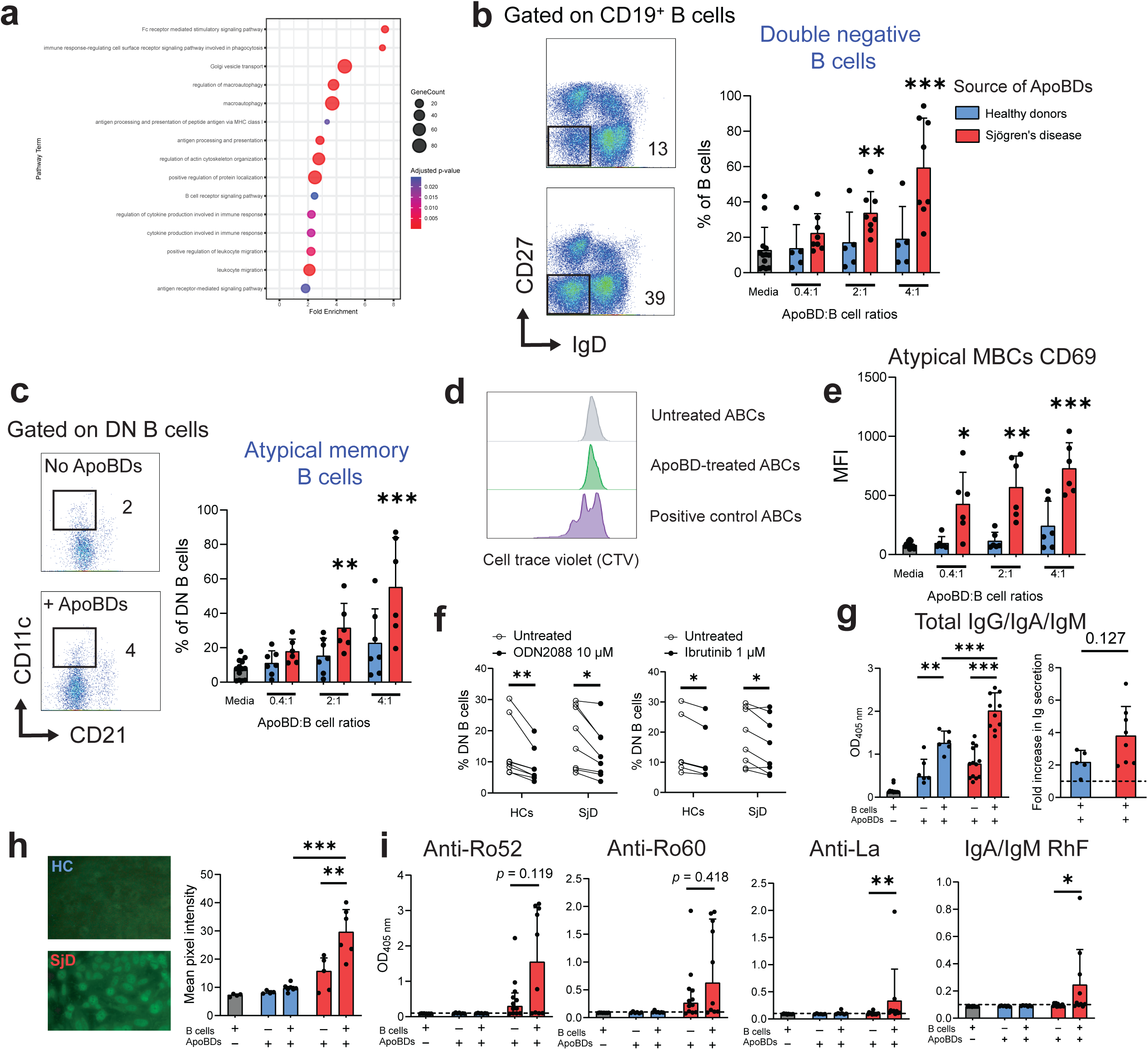
Apoptotic bodies (ApoBDs) cause the expansion and activation of atypical memory B cells. **(a)** Bubble plot of the top up-regulated immune-related biological processes of healthy donor (HC) B cells cultured with Sjögren’s disease (SjD) ApoBDs compared to unrelated HC ApoBDs, as assessed by mass spectrometry. ApoBDs from SjD or HC patients were co-cultured with purified B cells in a 4:1 ratio for 2 days. ApoBDs were removed and B cells assessed by mass spectrometry. Data is pooled from 3 SjD and 3 HCs ApoBDs cultured with B cells from 3 separate HCs from two independent experiments. **(b)** Proportion of double negative (DN) (IgD**^‒^**CD27**^‒^**) B cells after isolated peripheral B cells are cultured with ApoBDs from SjD and HC. Representative flow cytometry plots (left). The graph represents pooled data from 6 independent experiments. Pairwise comparisons against the untreated condition were performed by Friedman test with Benjamini-Hochberg FDR correction. **(c)** Proportion of atypical memory B cells (CD11c^+^ CD21^low^) gated from DN B cells. Representative flow cytometry plots (left). The graph represents pooled data from 6 independent experiments. Statistics were performed similarly as in (b). **(d)** Proliferation assay with atypical memory B cells. B cells were isolated from healthy donor PBMCs, labelled with cell trace violet (CTV) and co-cultured with or without ApoBDs in a 4:1 ratio for 48 hours. The plots are gated specifically on atypical memory B cells as in (c). Representative plot of two independent experiments with 2 healthy donor and 2 SjD-derived ApoBD samples. **(e)** CD69 mean fluorescence intensities (MFIs) measured on atypical memory B cells (MBC) from B cells cultured with HC and SjD ApoBD by flow cytometry. The graph represents pooled data from 6 independent experiments. Statistics were performed similarly as in (b). **(f)** Proportions of atypical memory B cells after treatment with 10 µM ODN2088 (Toll-like receptors 7-9 inhibitor) or 1 µM ibrutinib (Bruton’s tyrosine kinase inhibitor). Peripheral blood mononuclear cells from unrelated healthy donors were incubated with ApoBDs from HCs or SjD patients at a ratio of 4:1 (ApoBDs:B cell) with or without inhibitors. Cultures were harvested after 48 hours and subjected to flow cytometry. Pooled data from 5 independent experiments. Statistics were analysed using paired t tests with Šidák correction for multiple tests. **(g)** Antibody (IgG, IgA and/or IgM) secretion in B cell and ApoBD co-cultures measured by enzyme-linked immunosorbent assay (ELISA) (left). Supernatants diluted 1:10 and run in technical duplicates. Ratios were calculated for each ApoBD donor as OD (treated) / OD (untreated). Comparison on ratios were calculated with Mann-Whitney U test. **(h)** Indirect immunofluorescence microscopy of B cell-ApoBD culture supernatants on HEp-2 substrate (left). Neat supernatants were incubated on tissue slides followed by combined anti-human IgG/IgA/IgM FITC conjugate. Micrographs are representative images taken with 40x objective lens. Staining intensity was quantified using ImageJ software (right). Data are pooled from 3 independent experiments. **(i)** Specific antinuclear antibodies (ANAs) (anti-Ro52, anti-Ro60 and anti-La) and rheumatoid factors (RhF) measured in the B cell-ApoBD culture supernatants by ELISA. Neat supernatants were used and IgG/IgA/IgM conjugate used for ANAs, and IgA/IgM conjugate for RhF. Horizontal dotted lines represent cut-offs as determined by the mean + 2 standard deviations optical density (OD) value of 10 negative controls (media only). Data are pooled from 3 independent experiments.

Having identified an activated B cell protein signature upon exposure to SjD-ApoBD, we next assessed B cell subset composition after co-culture with SjD or healthy donor ApoBDs by flow cytometry (Supplementary Figure 2d). Addition of SjD-derived ApoBDs induced a dose–dependent increase of IgD^−^CD27^−^ double–negative (DN) memory B cells (Figure 2b) and atypical memory B cells (Figure 2c) compared to heathy donor ApoBDs. Atypical memory B cells were defined as previously described(34) IgD^−^CD27^−^CD21^lo^CD11c^+^ (Supplementary Figure 2e). In contrast, all other major peripheral B cell subsets, were reduced (Supplementary Figure 2f). CellTrace Violet labelling and Ki67 analysis revealed no evidence of DN or atypical memory B cell proliferation (Figure 2d and Supplementary Figure 2g), indicating these subsets may have differentiated from other B cell subsets rather than *in vivo* proliferation. Consistent with recent differentiation, atypical memory B cells also exhibited increased expression of the early activation marker CD69 (Figure 2e).

Given that atypical memory B cell differentiation depends on dual engagement of BCR and TLR7/9(35, 36), we surmised that autoantigens and nucleic acids in ApoBDs (Supplementary Figure 2h) stimulate BCR and TLR7/9 respectively(8, 37) to induce atypical memory B cell expansion. To test this, we assessed B cell subset composition after ApoBD co-culture with BCR and TLR7/9 inhibitors. To determine working concentrations of the inhibitors, DN B cell cultures were established as previously described (35), and ibrutinib and ODN2088 were titrated to achieve a ∼50% reduction in DN B cells without affecting viability (Supplementary Figures 2i, 2j). Pharmacological blockade of TLR7/9 (ODN2088) or BCR signalling (ibrutinib) significantly reduced atypical B cell expansion induced by both healthy control and SjD-derived ApoBDs by approximately 20-40% (Figure 2f). Transient exposure (removal of ApoBDs after 4 hours of co-culture) was sufficient to induce atypical B cell expansion at 48 h (Supplementary Figure 2k), and no evidence of direct ApoBD engulfment by B cells was observed (Supplementary Figure 2l). Collectively, these data indicate that ApoBDs promote atypical B cell differentiation via BCR– and TLR–dependent signalling rather than proliferation or direct engulfment.

Atypical memory B cells have properties consistent with pre-plasma cells enabling their rapid expansion to produce antibodies and autoantibodies (38). We therefore quantified immunoglobulin in the supernatants of ApoBD-B cell co-cultures. ApoBDs derived from both healthy donors and SjD patients induced antibody secretion (IgG, IgA, and IgM measured as a composite read-out) compared to separate cultures of B cells or ApoBD alone. However, antibody production was significantly more pronounced in B cell cultures exposed to SjD-derived ApoBDs (Figure 2g). Normalisation to donor specific ApoBD only controls revealed an approximately two-fold increase in immunoglobulin production following exposure to healthy control ApoBDs, compared with an approximately four-fold increase induced by SjD ApoBDs (Figure 2g). The supernatants from SjD-derived ApoBD-B cell co-cultures were enriched with antinuclear antibodies as determined using indirect immunofluorescence on HEp–2 substrates (Figure 2h). Specifically, increased levels of disease-relevant autoantibodies, anti-SSB/La and rheumatoid factor, were measured in the supernatants of B cells cultured with SjD ApoBDs but not heathy donor ApoBDs (Figure 2i). Together, our data indicates that SjD ApoBDs drive the expansion of atypical memory B cells and secretion of autoantibodies in vitro.

### Sjögren’s disease-derived apoptotic bodies induce inflammation in monocyte-derived macrophages in part via Toll-like receptors and Fc receptors

In addition to B cell hyper-activity, SjD is characterised by chronic innate immune activation and macrophage–driven inflammation (39, 40). Although apoptotic corpses in SjD render efferocytosis defective, phagocytes retained the capacity to engulf ApoBDs equivalent to healthy donors (Figure 1f). We therefore asked whether efferocytosis of SjD–derived ApoBDs qualitatively reprograms macrophages toward an inflammatory phenotype. To this end, healthy donor MDMs were co–cultured with ApoBDs from healthy controls or SjD patients. Following efferocytosis, MDMs were sorted into engulfing (CypHer5E⁺) and non–engulfing (CypHer5E⁻) populations and cultured for a further 18 hours. Culture supernatants revealed significantly increased secretion of IL–8 and CCL2 from MDMs that had engulfed SjD ApoBDs compared with healthy control ApoBDs (Figure 3a), whereas IL–10, IL–13, IL–1β, IFN–α and IFN–γ were not detected (Supplementary Figure 3a). Transcriptomic profiling of MDMs demonstrated up–regulation of immune–related gene programs, including antiviral defence and antigen presentation pathways, in SjD ApoBD–engulfing MDMs. The corresponding cytokine genes (Figure 3a) were not up-regulated (Figures 3b, 3c).

**Figure 3.**
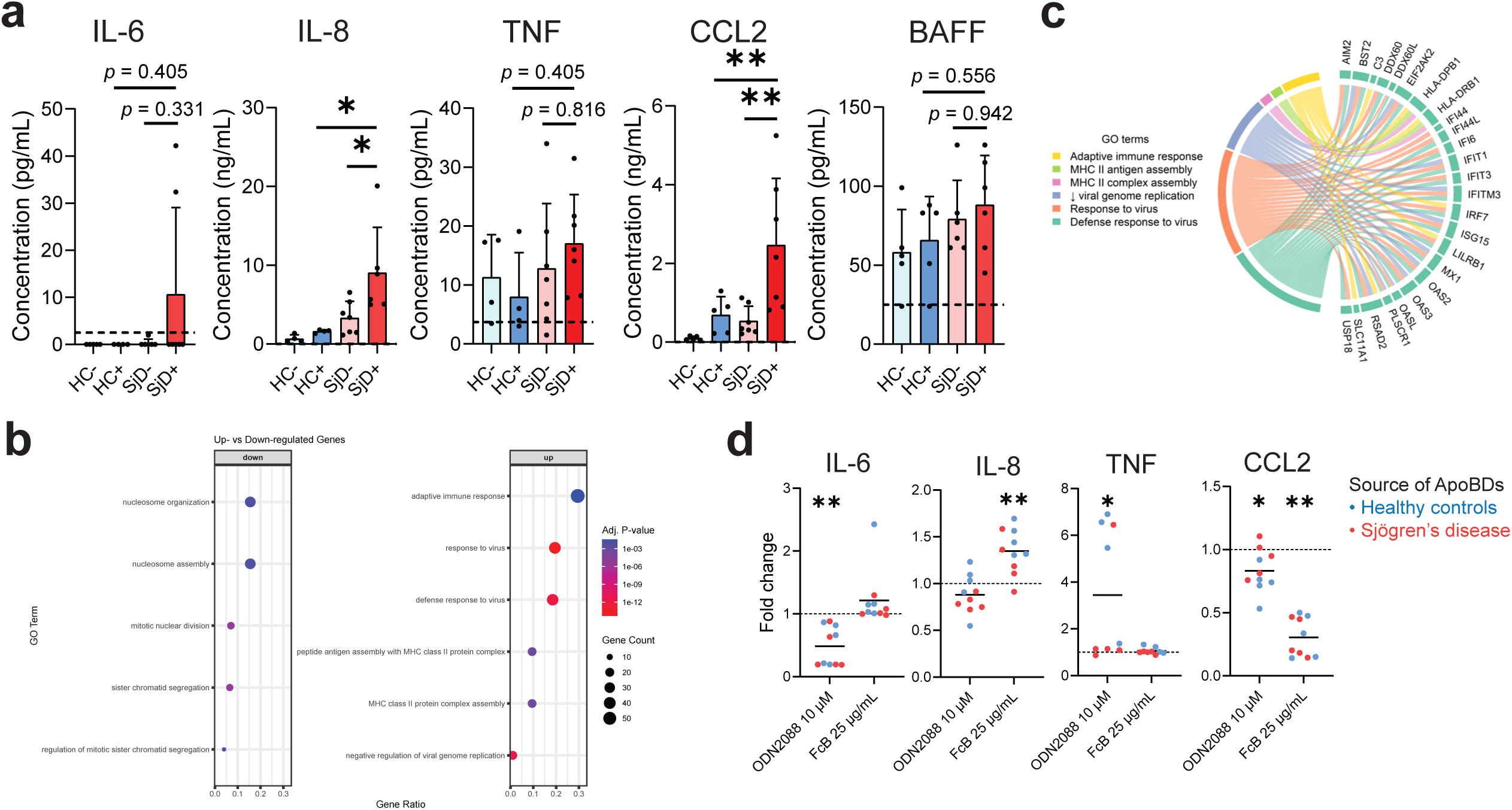
Sjögren’s disease (SjD)-derived apoptotic bodies (ApoBDs) are inflammatory and cause monocyte-derived macrophage (MDM) activation in vitro. **(a)** Cytokines in the supernatant from MDM cultures measured by cytokine bead array (flow cytometry). HC/SjD^−^ represents MDMs that did not engulf HC/SjD ApoBDs; HC/SjD^+^ are MDMs that engulfed HC/SjD ApoBDs. Columns represent mean with standard deviations. *TNF*, tumour necrosis factor. *CCL2*, CC chemokine ligand 2. *BAFF*, B cell activating factor. **(b)** Gene ontology pathway analyses of top 100 differentially-expressed genes down- and up-regulated in MDMs engulfing SjD ApoBDs vs. healthy donor ApoBDs. A total of 5 samples for each condition were analysed and pooled. **(c)** Chord diagram showing up-regulated genes in monocyte-derived macrophages (MDMs) that have engulfed SjD ApoBDs) vs. those that have engulfed healthy donor ApoBDs. **(d)** Fold reduction of cytokines produced by MDM cultures following addition of ApoBDs from HC or SjD patients (untreated), and with or without inhibitors (ODN2088 or FcγR blockers [FcB]). Fold changes were tested against a null of no change (fold change = 1.0) by Wilcoxon signed-rank test with Benjamini-Hochberg correction.

Given that SjD ApoBDs contain nucleic acids and increased immunoglobulins compared to healthy donor ApoBDs, we next tested whether inflammatory activation of macrophages was mediated through TLRs and/or Fcγ receptors (FcγRs). Healthy donor MDMs were pre–treated with a TLR7/9 antagonist (ODN2088) or FcγR–blocking antibody prior to ApoBD exposure. Both pharmacological inhibitors significantly reduced CCL2 secretion in response to ApoBDs, while IL-6 secretion was only reduced by the TLR7/9 antagonist. TNF levels were unaffected and FcγR blockade unexpectedly enhanced IL–8 production (Figure 3d). Treatment with inhibitors alone had no effect on basal cytokine secretion (Supplementary Figure 3b). Collectively, these findings demonstrate that, despite impaired efferocytosis of SjD ApoBD, engulfment of patient–derived ApoBDs actively drives inflammatory programming in healthy donor-derived MDMs through TLR– and FcγR–dependent pathways, thereby converting a normally tolerogenic process into a pro–inflammatory response.

### Apoptotic bodies require nuclear cargo and inflammatory environment to drive atypical memory B cell and macrophage activation

Having established that SjD-derived ApoBDs are intrinsically capable of reducing efferocytosis, driving atypical B cell expansion and inflammatory macrophage activation, we next compared the molecular determinants of ApoBDs from SjD and healthy donors to identify factors that associate with their conversion from tolerogenic substrates into immunostimulatory signals.

To obtain an unbiased overview of ApoBD composition, we performed mass spectrometry-based proteomic profiling of ApoBDs isolated from pooled plasma samples of SjD patients and healthy donors. The majority of proteins detected in ApoBDs localised to the cytoplasm and nucleus (Figure 4a) and were associated with metabolic and cell communication pathways (Figure 4b). Comparative analysis identified multiple differentially expressed proteins in SjD-derived ApoBDs relative to healthy controls (Figure 4c). Notably, SjD ApoBDs exhibited significant enrichment of proteins involved in cellular metabolism (Supplementary Figure 3c), and inflammation such as APOE, C5, IRAK4 and JAK2 (Figure 4d), consistent with heightened immune activation in SjD. Given the central role of nuclear antigens in systemic autoimmunity, we next examined the abundance of disease-relevant autoantigens within ApoBDs (41, 42). SjD-associated autoantigens, including Ro52/TRIM21, Ro60/TROVE2 and other ribonucleoproteins (42, 43), were present at higher levels in SjD-derived ApoBDs compared with healthy controls (Figure 4e). In contrast, histone proteins, also common autoimmune targets, were reduced in SjD ApoBDs (Figure 4e). Taken together, these analyses indicate that the defining feature of SjD ApoBDs is a selectively amplified inflammatory signature, likely primed from an *in vivo* inflammatory environment of autoimmunity.

**Figure 4.**
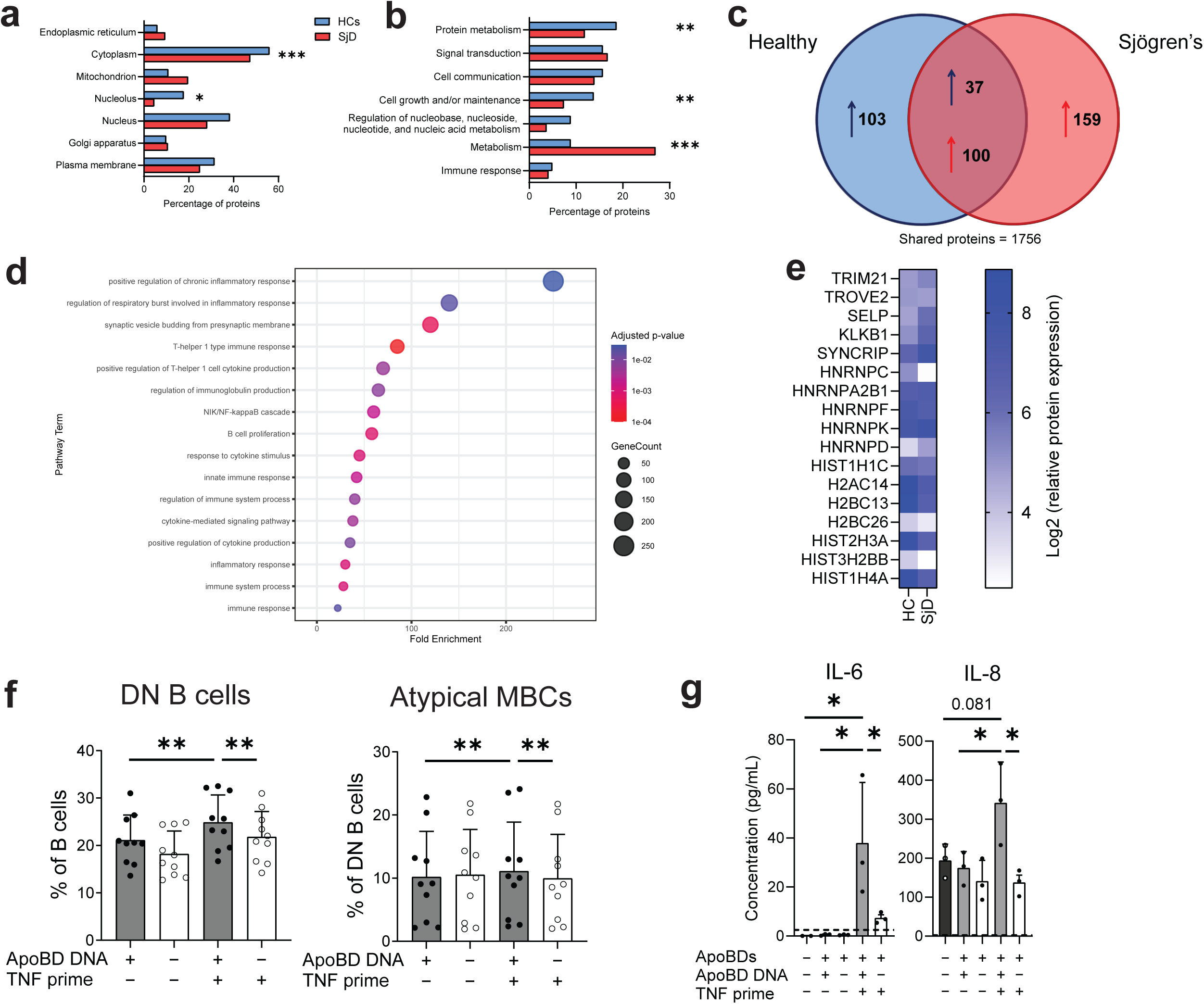
Sjögren’s disease apoptotic body (ApoBD) contents contribute to B cell and macrophage (MDM) activation. **(a)** Proportions of proteins, assessed by mass spectrometry, attributed to cellular location and identified in healthy donor (HC) and Sjögren’s disease (SjD) ApoBDs according to the UniProt Homo Sapiens database. **(b)** The proportion of SjD and HC ApoBD proteins with biological functions according to the UniProt Homo Sapiens database. **(c)** Venn diagram of differentially expressed proteins in HC and SjD ApoBDs, as detected by mass spectrometry. One hundred and three proteins are upregulated and present only in HCs, while 159 proteins are only present in SjD patients and are up-regulated. Within the shared protein group, 37 proteins are upregulated in HCs, and 100 proteins are upregulated in SjD. **(d)** Gene ontology pathway enrichment analysis of the top up-regulated immune system-related proteins in SjD ApoBDs relative to HC ApoBDs. **(e)** Autoantigen expression heat map for ApoBDs derived from HC and SjD patients. **(f)** Percentage and mean fluorescence intensity (MFI) ratios of double negative (DN) B cells and atypical memory B cells (MBCs) following healthy donor B cell co-culture with Jurkat cells. ApoBD DNA+ represents Jurkat cell ApoBDs that are DNA replete (sgPANX1 cell line) while ApoDNA– represents Jurkat cell ApoBDs that are DNA deficient (double knock-out for caspase-activated DNase [CAD] and pannexin 1). For TNF priming, Jurkat cells were incubated with TNF 500 ng/mL 24 hours before apoptosis induction to create ApoBDs. Each dot represents a separate B cell healthy donor. Data is pooled from two independent experiments. Paired comparisons were performed by Wilcoxon signed-rank tests with Benjamini-Hochberg adjustments. **(g)** Cytokines in supernatant from MDM co-cultures with Jurkat cell ApoBDs. Data is pooled from two independent experiments with 3 healthy donor MDMs.

To disentangle the roles of nucleic acid cargo and an inflammatory environment driving ApoBD-mediated B cell activation and macrophage inflammation, we generated a nuclear-depleted ApoBD model through genetic disruption of caspase-activated DNase (CAD) in Jurkat T cells. We also manipulated the packaging of inflammatory cargoes into these ApoBDs using TNF priming during apoptosis induction. Following an ApoBD-B cell co-culture, inflammatory, nuclear content-carrying ApoBDs induced the most robust expansion and activation of DN and atypical memory B cells. By contrast, those lacking either inflammatory priming or nuclear materials attenuated this effect. Although the absolute difference between conditions was modest for atypical memory B cells, the change was highly consistent across subjects (9/10 donors, Wilcoxon Benjamini-Hochberg-adjusted p=0.006), reflecting a reproducible within-subject effect (Figure 4f). Correspondingly, MDMs engulfing nuclear-carrying, inflammation-primed ApoBDs mounted robust inflammatory cytokine responses upon efferocytosis, whereas DNA-depleted or non-inflammatory ApoBDs failed to secrete pro-inflammatory cytokines (Figure 4g). Together, these data demonstrate that the capacity of ApoBDs to drive pathogenic B cell differentiation and macrophage inflammation is dictated by their nuclear cargo and inflammatory context, establishing ApoBDs as active conveyors of immunostimulatory signals in SjD rather than inert by-products of cell death.

## Discussion

Apoptosis and its resolution through efferocytosis are fundamental processes for maintaining immune tolerance, and defects in apoptotic cell clearance have long been implicated in systemic autoimmune diseases (44). Yet how the accumulated ApoBDs circulating in patients with SjD contributes to disease pathogenesis has not been fully defined. In this study, we establish qualitatively altered ApoBDs as a previously unrecognised intrinsic driver of defective efferocytosis and immune dysregulation in SjD, providing a mechanistic link between impaired cell death clearance and both adaptive and innate immune activation.

We identify a marked increase in circulating ApoBDs in patients with SjD that reflects defective efferocytosis rather than an increased apoptotic burden or intrinsic phagocyte dysfunction. Although apoptosis is elevated in SjD, MDMs retained the capacity to engulf ApoBDs in vitro, indicating that impaired clearance is not explained by saturation of phagocytic capacity alone. Instead, SjD–derived ApoBDs exhibited reduced efferocytic uptake, a defect that was partly recapitulated when ApoBDs were coated with patient plasma. These findings support previous findings that SjD-associated autoantibodies can contribute to defective efferocytosis (14, 17, 18) thereby allowing immunostimulatory ApoBDs to accumulate in circulation.

The accumulation of ApoBDs in SjD has direct functional consequences on both adaptive and innate immune compartments. ApoBDs promoted the expansion of atypical memory B cells and *de novo* autoantibody secretion, consistent with previous findings that CD21^low^ atypical memory B cells are enriched for autoreactivity and poised for plasma cell differentiation (32, 45). In parallel, efferocytosis of SjD ApoBDs actively reprogrammed MDMs toward an inflammatory phenotype instead of canonical immune tolerance, thus potentially contributing to SjD-associated chronic inflammation. While apoptotic material has previously been shown to activate innate immune cells under certain contexts (46), our study demonstrates that patient–derived ApoBDs are sufficient to elicit these responses *ex vivo*, identifying ApoBDs as active mediators rather than passive markers of defective efferocytosis.

A major mechanistic insight from this study lies in defining the ApoBD features that confer pathogenic potential. Unbiased proteomic profiling revealed heterogeneous changes across metabolic and nuclear proteins between SjD-derived and healthy control ApoBDs; notably, the most prominent and consistent distinguishing feature was the selective enrichment of inflammation–associated proteins in SjD ApoBDs. This inflammatory signature emerged as the defining characteristic of disease–derived ApoBDs. When coupled with nuclear materials, which have long being recognised as a dominant driver of autoimmune manifestations (3), this inflammatory context endows SjD ApoBDs with the capacity to elicit robust B–cell expansion and immune activation. Together, these findings support a model in which ApoBDs function as composite danger signals, whereby nuclear autoantigens engage B–cell receptors, nucleic acids activate endosomal Toll–like receptors, and antibody coating facilitates Fcγ receptor signalling, with these inputs acting in concert to amplify pathogenic immune responses rather than any single ligand operating in isolation.

We therefore propose a model in which defective efferocytosis in SjD permits the accumulation of inflammatory, nuclear–laden ApoBDs that act as central hubs coordinating B cell autoreactivity and MDM inflammation. By converting a normally tolerogenic process into a sustained source of immune activation, these altered ApoBDs provide a mechanistic framework linking impaired apoptotic clearance to chronic autoimmunity in SjD. A limitation of the present study is its reliance on *ex vivo* human systems, which precludes direct assessment of tissue–specific effects within exocrine glands and other organs of interest. Nevertheless, our findings establish a conceptual foundation for future *in vivo* studies and suggest that restoring ApoBD clearance or disrupting ApoBD–driven immune sensing may represent therapeutic strategies with relevance beyond SjD.

## Methods

### Patients

Whole blood was obtained from SjD patients or healthy donors into ethylenediaminetetraacetic acid tubes by venepuncture. SjD patients met the American College of Rheumatology/European Alliance of Associations for Rheumatology classification criteria for primary SjD (47), and healthy donors were derived from consenting subjects with no known chronic or autoimmune diseases. Healthy donor PBMCs were obtained from anonymised blood donations from the Australian Lifeblood Service. ApoBDs were obtained from whole blood and processed fresh within 2 hours of collection. Isolation was performed by centrifugation as previously described (28). Briefly, whole blood was centrifuged 200 × *g* for 15 min, plasma removed and then centrifuged at 500 × *g* for 10 min. The supernatant was centrifuged at 3000 × *g* for 20 min, and the pellet washed twice with PBS. ApoBD purity was consistently >97%. All patients provided written, informed consent for participation. Studies were approved by the Western Sydney Local Health District Research Office (ETH01030).

### Flow cytometry and fluorescence-activated cell sorting (FACS)

Whole blood underwent red blood cell lysis using ammonium chloride solution (Kinetik, Australia). PBMCs or whole blood were stained in cocktails of antibodies from BD Biosciences: CD3 APC-H7 (SK7), CD10 BUV737 (HI10a), CD10 PE-CF594 (HI10a), CD11c BUV395 (B-ly6), CD14 BV650 (clone M5E2), CD19 BV605 (clone SJ25C1), CD45 BV786 (HI30), CD56 PE-Cy7 (NCAM16.2) and/or CD303 APC (AC144, Miltenyi Biotec).

ApoBDs and apoptotic cells were identified flow cytometrically with the following markers: annexin V-BV421, propidium iodide (#P1304MP, Invitrogen), FLICA (#I35102, Invitrogen) and/or TO-PRO3 iodide (#T3605, Invitrogen) as previously described.(28) Fixation and permeabilisation of ApoBDs was performed using Cytofix/Cytoperm fixation and permeabilisation solution (#554722, BD) or transcription factor buffer set (#562574, BD). Samples were acquired on an LSRFortessa cytometer (BD) or sorted via a FACSAria cell sorter (BD).

### Mass spectrometry

For ApoBD proteomics, liquid chromatography-mass spectrometric (LC-MS) analysis was performed on an Orbitrap Eclipse Mass Spectrometer (ThermoFisher Scientific). The liquid chromatography systems were equipped with an Acclaim Pepmap nano-trap column (Dinoex-C18, 100 Å, 75 µm x 2 cm) and an Acclaim Pepmap RSLC analytical column (Dinoex-C18, 100 Å, 75 µm x 50 cm). Data-independent acquisition (DIA) data was analysed using Spectronaut software (v. 17.5.230413.55965) against the UniProt *Homo Sapiens* database (updated September 2023) using the default search parameters. The false discovery rate (FDR) for proteins and peptide-spectrum match for both Pulsar and DIA analysis was set to 1%. Bioinformatic analyses of proteins were based on label-free quantitation of 3 healthy donor and 3 SjD ApoBD samples as previously described.(48) Student’s t test was performed to look for differences between means, with P values permutation-based and FDR-adjusted for a threshold of 5%.

For B cell culture proteomics, mass spectrometry was performed via the Biomedical Proteomics Facility at the Children’s Medical Research Institute (Westmead, Australia). Frozen pellets of B cells were digested with trypsin before running on a Vanquish Neo UHPLC system and Astral Orbitrap mass spectrometer (Thermo Fisher Scientific). The raw LC-MS/MS data was processed with DIA-NN v2.2. The *Homo sapiens* reference proteome (using one gene per protein) downloaded Sep 30 2025 from UniProtKB with 20,663 genes was used to create an *in silico* library.

### Monocyte-derived macrophage cultures and engulfment assays

Type 1 MDM cultures were established from fresh human buffy coats as similarly described (49). Briefly, monocytes were seeded onto a 6- or 48-well plate at a density of 2.0 x 10^6^/well or 0.2 x 10^6^/well respectively and cultured for at least 6 days. As a source of ApoBDs for engulfment assay, HEK293 cells (Expi293F cell lines, Gibco) seeded into 6-well plates at a concentration of 0.5 x 10^6^ cells/mL and apoptosis induced using 2 µM of raptinal (#HY-121320, MedChemExpress) for 3 hours, 37°C and 5% CO_2_. Cytometric bead arrays for cytokines (BD Biosciences) were performed on neat supernatants in technical duplicates for interleukin (IL)-1β, IL-6, IL-8, IL-10, IL-13, IFN-α, IFN-γ, CC chemokine ligand (CCL) 2 and TNF. ApoBDs derived from patients were incubated with MDMs for 1.5 hours to examine efferocytosis. For FACS, MDMs were stained CypHer5e dye and sorted into engulfers (CypHer5e^+^) or non-engulfers (CypHer5e^−^) as previously described (49). For bulk total RNA sequencing, total RNA was extracted for harvested MDMs via the RNeasy Micro Kit (Qiagen) before library preparation with SMART-Seq Total RNA Library Prep (Takara Bio).

The TLR7-9 inhibitor, ODN2088 (#130-105-816, Miltenyi Biotec) and Fcγ receptor blocker (#564220, BD) were used at 10 µM and 25 µg/mL as per manufacturer’s instructions. For microscopy, MDMs were plated on microscopy chambers and examined using the Stellaris Confocal Microscope platform (Leica).

### Jurkat cell cultures

PANX1^−/−^ Jurkat T cells were generated using CRISPR-Cas9 gene editing as per previously described (37). PANX1^−/−^ CAD^−/−^ Jurkat T cells were obtained by electroporating pX330 vector encoding sgRNA (50) into PANX1^−/−^ cells and limiting dilution. CAD deficiency was confirmed by western blotting and DNA fragmentation assay following apoptotic stimuli. DNA-enriched or DNA-deplete ApoBDs were generated by inducing apoptosis in the PANX1^−/−^ or PANX1^−/−^ CAD^−/−^ respectively, using UV radiation or 1 µM of raptinal. These ApoBDs were cultured with MDMs or B cells at a ratio of ApoBDs:target cell of 3:1.

### B cell cultures

B cells were isolated from cryopreserved HC PBMCs by negative selection using the human pan B cell isolation kit (#130-101-638, Miltenyi Biotec) and plated into 48-well plates at a density of 1 x 10^6^/mL of B cell media. The media comprised RPMI-1640 media, 10% foetal calf serum, 55 µM of β-mercaptoethanol, 1x GlutaMAX (Gibco), penicillin-streptomycin 100 U/mL and 100 µg/mL, 1 mM sodium pyruvate (Gibco) and 10 mM HEPES (Gibco). B cells were cultured with or without ApoBDs derived from HCs or SjD patients for 48 hours at 37°C and 5% CO_2_. The TLR 7, 8 and 9 inhibitor, ODN2088 (#130-105-816, Miltenyi Biotec), B cell receptor inhibitor, ibrutinib (#HY-10997, MedChemExpress), RNase A (#R5503, Sigma-Aldrich) or DNase I (#11284932001, Roche) were co-cultured with B cells with and without ApoBDs. Following culture, the B cells were harvested for flow cytometric analyses and the supernatants snap frozen for antibody and cytokine analyses. For proteomic analyses, the B cell pellet was washed twice in PBS before being immediately frozen at −80°C. The purity of B cell cultures was consistently >95%.

### Antibody / autoantibody assessments

Enzyme-linked immunosorbent assays (ELISAs) were performed on B cell culture supernatants as previously described (51). ELISA plates were coated using 0.5 µg/mL of anti-human IgG (#2015-01, SouthernBiotech), anti-IgA (#2050-01, SouthernBiotech), anti-IgM (#109-005-043, Jackson); 1 µg/mL of Ro52 antigen (#ATR05), Ro60 (#ATR02), La (#ALA01, all Arotec Diagnostics) or 5 µg/mL Fc fragment (#ab90285, Abcam) as appropriate. Antinuclear antibody assessments were made by using commercial HEp2 slides (HEp-20-10, #FA 1522-2005, Euroimmun) and an anti-IgG/IgA/IgM FITC conjugate (#A18848, Invitrogen).

### Statistics

Statistical analyses were conducted using GraphPad Prism software. Mean values with standard deviation bars are plotted on column graphs. The specific statistical tests used in the analyses are listed in the respective figure legends. Summary statistic asterisks are as follows: * *p* < 0.05. ** *p* < 0.01. *** *p* < 0.001.

## Acknowledgements

The authors gratefully acknowledge the contributions of the patients who donated their samples for this research. In addition, the authors wish to thank Professors Mark Hulett and Ivan Poon (La Trobe University) for their insightful suggestions. ELISAs, flow cytometry, imaging and RNA sequencing were performed at the Westmead Scientific Platforms, which are supported by the Westmead Research Hub, the Westmead Institute for Medical Research, the Cancer Institute New South Wales, the Medical Research Future Fund, the National Health and Medical Research Council and the Ian Potter Foundation. Additional imaging and flow cytometry/cytokine data acquisition were performed at the La Trobe Bioimaging Platform (La Trobe University).

## Author contributions

Conceptualization: A.Y.S.L., M.W.L., T.K.P., J.H.R.

Formal analysis: A.Y.S.L., B.S., S.G., P.F., T-K.M.T., Q.T.L., T.K.N., T.K.P.

Investigation: A.Y.S.L., B.S., S.G., P.F., T-K.M.T., Q.T.L., C-S.A., T.K.N., T.K.P., J.H.R.

Resources: A.Y.S.L., M.W.L., T.K.P.

Data curation: A.Y.S.L., B.S., S.G., P.F., T-K.M.T., Q.T.L., C-S.A., T.K.N., T.K.P., J.H.R.

Writing – original draft: A.Y.S.L., T.K.P., J.H.R. Supervision: T.K.P., J.H.R.

Funding acquisition: A.Y.S.L., T.K.P., J.H.R.

All authors have critically read and revised the manuscript for important intellectual content.

## Data availability

The data that support the findings for this paper are available from the corresponding authors upon reasonable request. The mass spectrometry proteomics data have been deposited to the MassIVE repository (Center for Computational Mass Spectrometry, UCSD) under the dataset identifier MSV000101473.

