## Supplementary Figure for "Apoptotic bodies from patients with Sjögren’s disease drive atypical memory B cell and macrophage activation and autoantibody production"

### Supplementary Figure 1

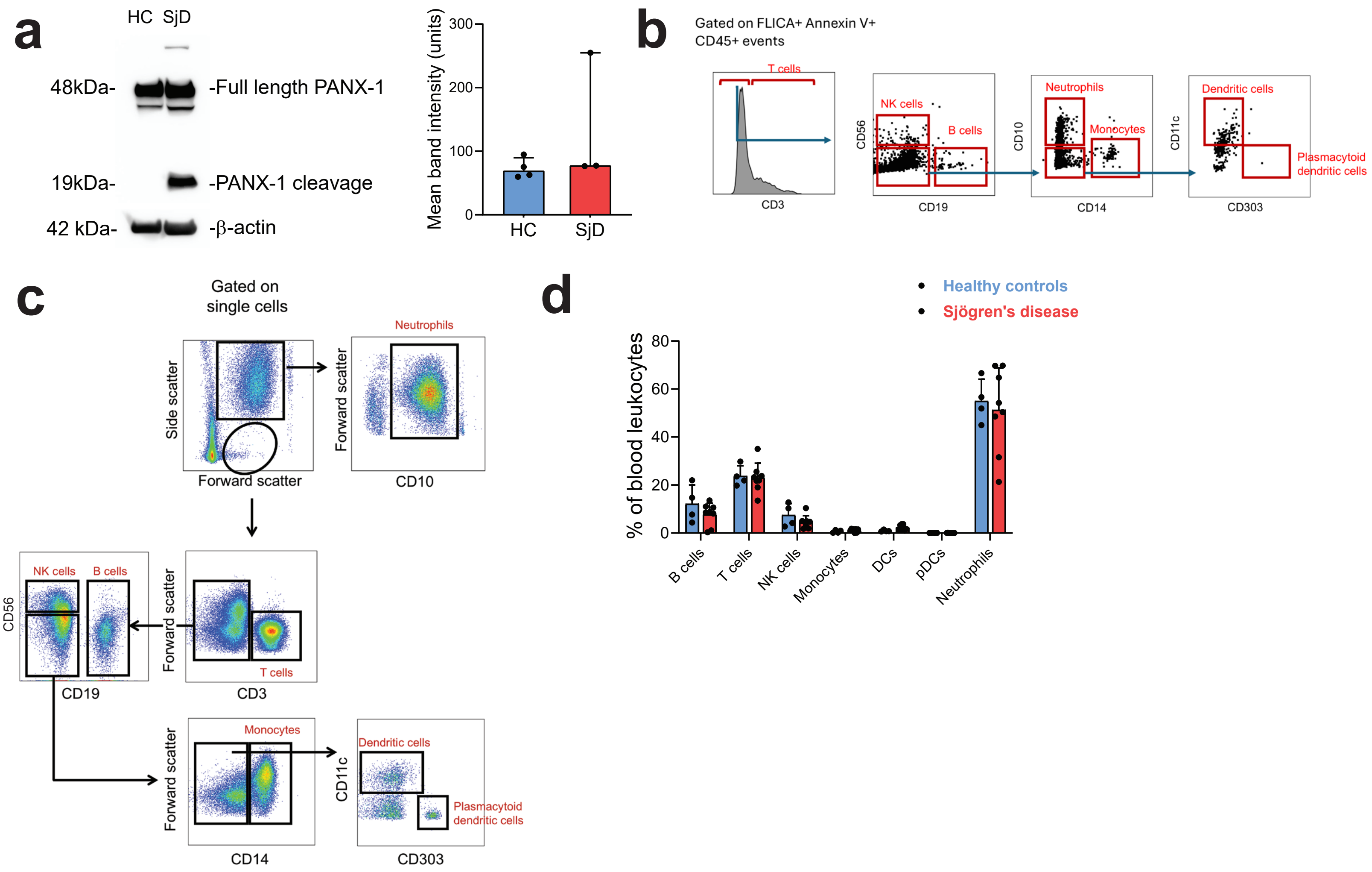


**(a)** Pannexin-1 (PANX-1) expression and cleavage products as analysed by Western blot. ImageJ was used to quantify the band intensities in the 3 Sjögren's disease (SjD) patients and 4 healthy donors (HCs) on an over-exposed gel. **(b)** Gating strategy for immunophenotyping CD45+ apoptotic bodies from Sjögren's disease (SjD) and healthy donors (HCs). The major immune cell populations were defined as follows: T cells (CD3+), natural killer (NK) cells as CD3−CD56+CD19−, B cells as CD3−CD56−CD19+, neutrophils as CD10+CD14− events negative for the aforementioned lineage markers, monocytes as CD10−CD14+ events negative for the aforementioned lineage markers, dendritic cells (DCs) as CD11cbrightCD303− and negative for the aforementioned lineage markers, and plasmacytoid dendritic cells (pDCs) as CD11c−CD303+ and negative for the aforementioned lineage markers. **(c)** Gating strategy and definitions of the peripheral blood immune cells. **(d)** Proportion of circulating leukocytes in blood of HCs and SjD as determined by flow cytometry. Each dot represents one patient, and percentages are expressed as a proportion of leukocytes for each of the indicated 7 populations for each patient. There were no statistical differences in the proportions across the patient groups (HC vs. SjD).

### Supplementary Figure 2

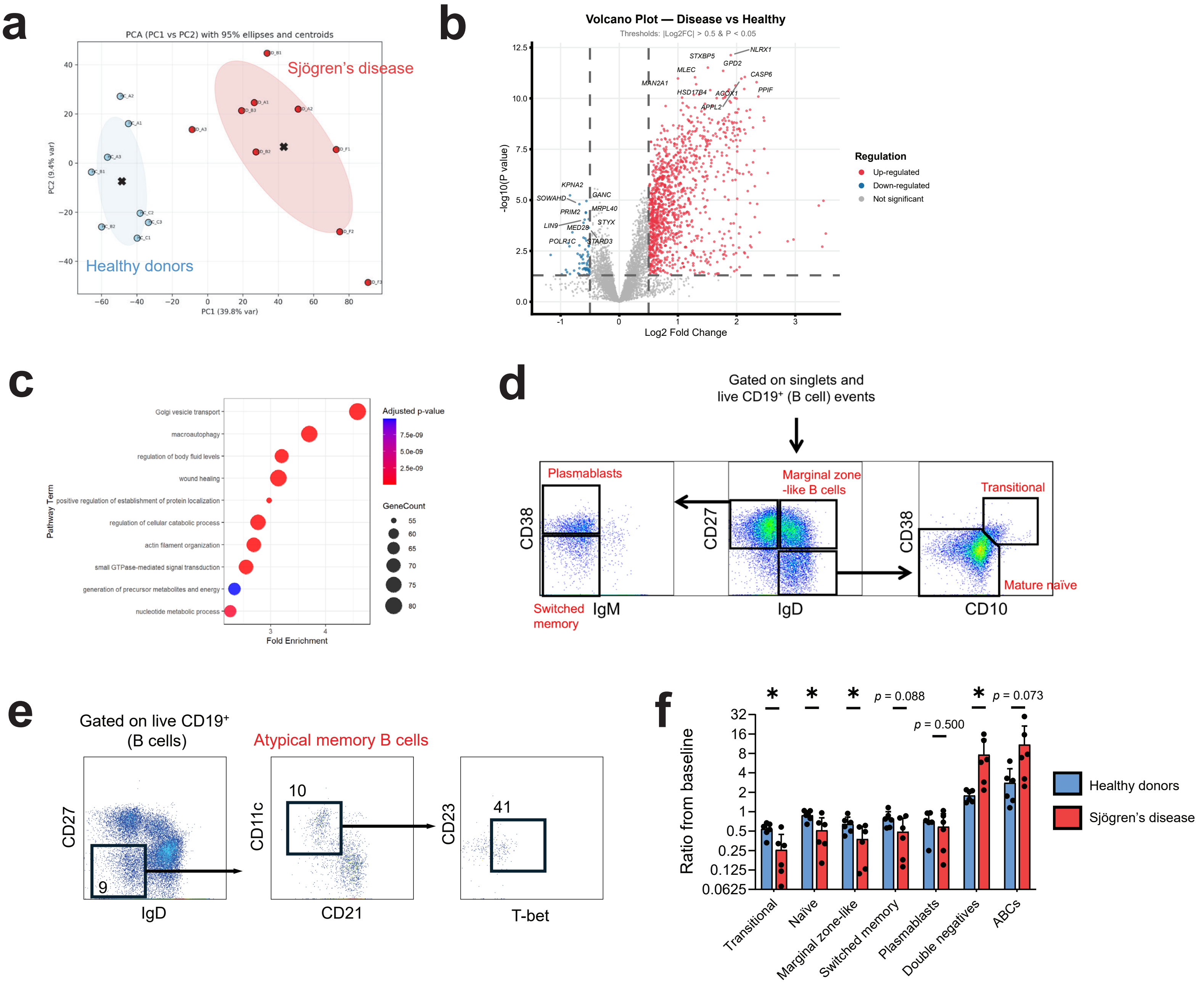


**(a)** Principal component analysis plot with 95% ellipses of B cell-ApoBD cultures. Red dots represent ApoBD from Sjögren's disease (SjD) patients whilst ApoBDs from healthy control (HCs) are in blue. Each dot represents a single culture, with those with the same letter representing an individual replicate of the same ApoBD donor. **(b)** Volcano plot of up-regulated and down-regulated proteins from mass spectrometric analyses of B cells co-cultured with apoptotic bodies (ApoBDs) derived from HCs (3 donors) or SjD (3 donors) at a ratio of 4:1 ApoBDs:B cell. B cells were spun at 300 × g and washed to isolate B cells from ApoBDs following the culture. Plot is representative of two independent experiments. The top 10 up- and down-regulated genes are marked. **(c)** Bubble plot demonstrating top ten up-regulated gene ontology biological pathways in B cells treated with SjD ApoBDs relative to HC ApoBDs. Mass spectrometry data is pooled for 3 HCs and 3 SjD patients. **(d)** Flow cytometric gating strategies to define B cell subsets. **(e)** Gating strategy for flow cytometric definition of atypical memory B cells. B cells from healthy donors were isolated by negative magnetic sorting. Atypical memory B cells were defined as live CD19+ events, IgD−CD27−CD21loCD11c+ B cells and were further validated as CD23lo and positive for the canonical transcription factor, T-bet. The T-bet+ gate was defined as per a fluorescence minus one (FMO) control. **(f)** Ratio of each major B cell subset, normalised to untreated controls, in B cell cultures after being treated with HC or SjD ApoBDs at a ratio of 4:1. B cells were collected after 48 hours co-culture and assessed by flow cytometry. Data is representative of 3 independent experiments.

### Supplementary Figure 2 continued

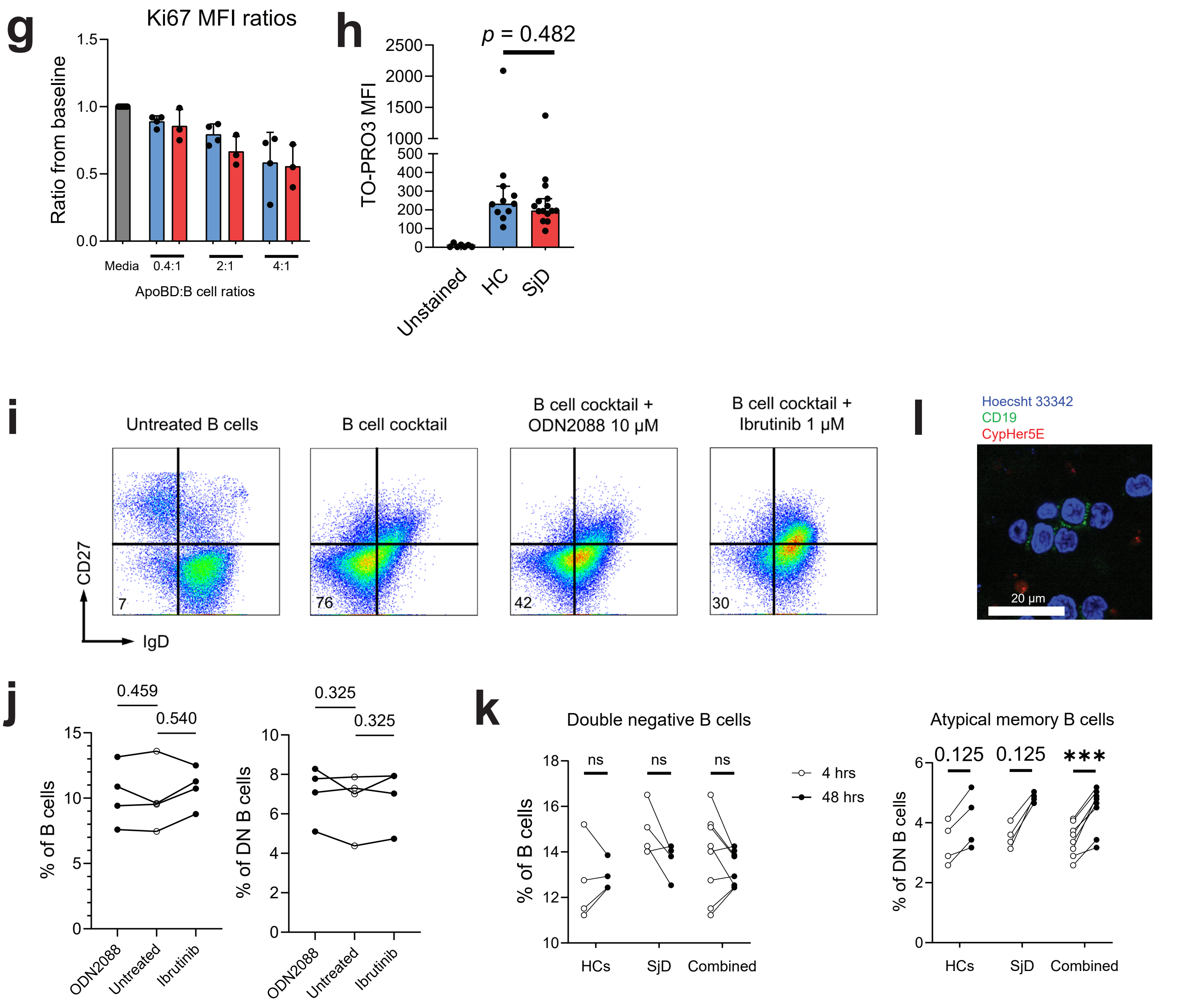


**(g)** Ki67 mean fluorescence intensity (MFI) ratios of atypical memory B cells from ApoBD and B cell co-cultures, normalised to a media-only control. Each dot represents either a single culture of ApoBDs derived from healthy donor or SjD patient. **(h)** Nucleic acid detection on HC and SjD ApoBDs detected by the nucleic acid-binding dye, thiazole orange-propidium 3 (TO-PRO3). ApoBDs were assessed flow cytometrically and TO-PRO3 assessed by its MFI. **(i)** Atypical memory B cell cocktail (see Methods) treatment of isolated B cells and impact of B cell inhibitors. B cells were cultured for a total of 2 days and plots are gated on total single and live B cells (CD19+ events). **(j)** ODN2088 and ibrutinib do not impact on the survivability of B cells. Isolated B cells were treated or not with ODN2088 10 µM or ibrutinib 1µM and the percentages of double negative (DN) or atypical memory B cells (of DN B cells) were compared to corresponding untreated samples by paired t tests. Cultures were left for 48 hours. A total of 4 healthy donors were used for this experiment. **(k)** Percentages of double negative (DN) B cells and atypical memory B cells after ApoBDs have been co-cultured for 4 hours (open circles) or 48 hours (black circles). For the 4-hour co-cultures, B cells were harvested from culture, ApoBDs separated by centrifugation and B cells suspended in fresh media for the remaining 44 hours. Wilcoxon signed-rank test with Benjamini-Hochberg correction was used to analyse paired tests. ns, not statistically significant. **(l)** Confocal microscopy of B cell-ApoBD co-cultures. ApoBDs stained with CypHer5E and B cells isolated from healthy donors and then stained with CD19-AF488. The micrograph was taken with 63x objective lens and is a representative image of 4 independent experiments with ApoBDs derived from HCs (3) and SjD patients (3).

### Supplementary Figure 3

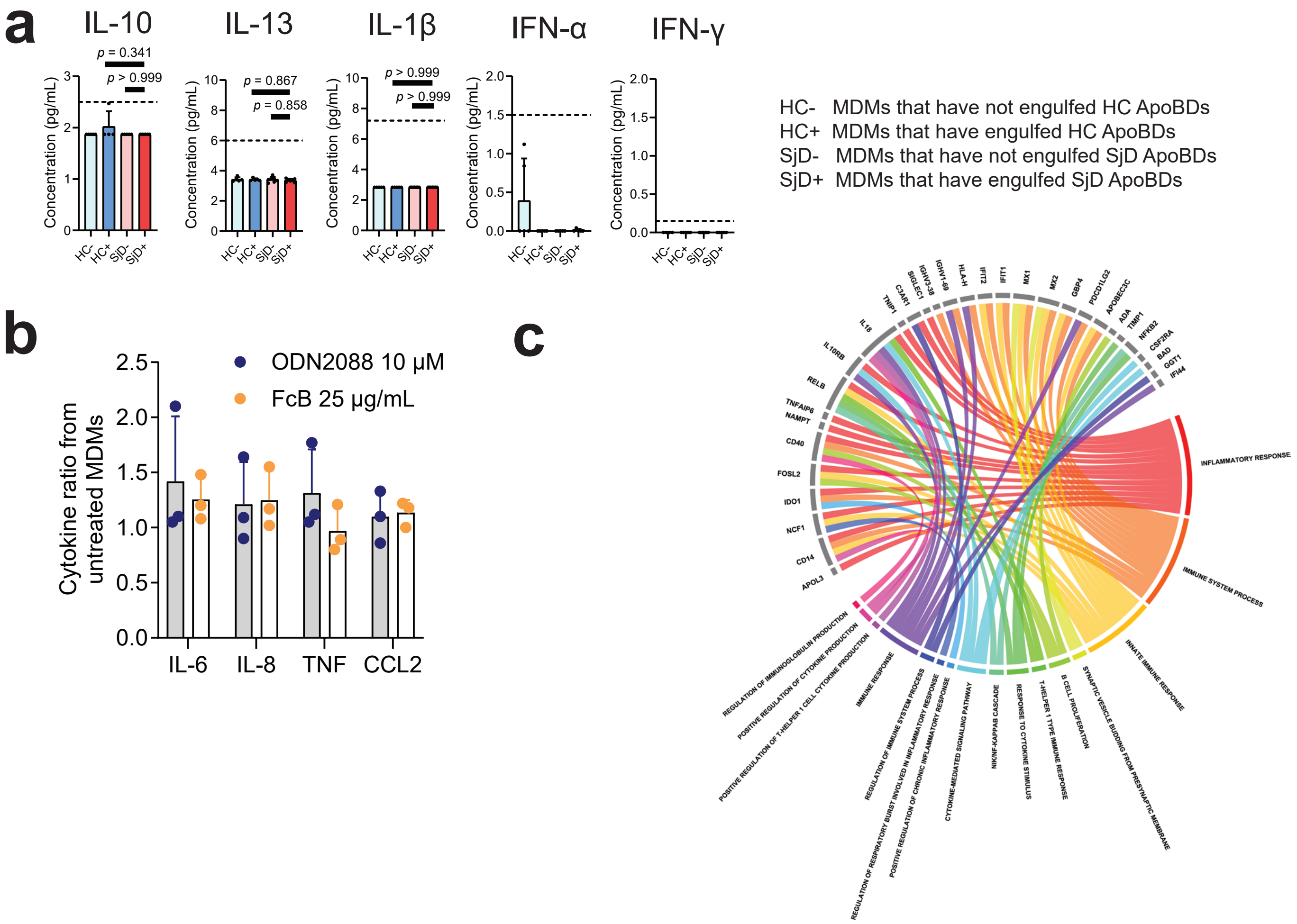


**(a)** Supernatant IL-10, IL-13, IL-1β, IFN-α and IFN-γ from healthy donor (HC) monocyte-derived macrophage (MDM) cultures with ApoBDs from either HCs or Sjögren's disease (SjD) patients. Cytokines were measured using the cytometric bead array cytokine kits (BD). Horizontal dotted lines represent the assay's limit of detection. Data were analysed using one-way ANOVA with Tukey's correction for multiple comparisons. **(b)** Ratios of IL-6, IL-8, TNF and CCL2 concentration in MDM cultures incubated with ODN2088 or Fcγ receptor blocker (FcB) compared to untreated MDMs. **(c)** Chord diagram showing up-regulated immune system-related proteins in SjD ApoBDs relative to healthy donor ApoBDs.
